# Suitability Evaluation of Toehold Switch and EXPAR for Cell-Free MicroRNA Biosensor Development

**DOI:** 10.1101/2023.05.11.540462

**Authors:** Caroline E. Copeland, Yong-Chan Kwon

## Abstract

The development of a robust and cost-effective sensing platform for microRNA (miRNA) is of paramount importance in detecting and monitoring various diseases. Current miRNA detection methods are marred by low accuracy, high cost, and instability. The toehold switch riboregulator has shown promising results in detecting viral RNAs integrated with the cell-free system (CFS). This study aimed to leverage the toehold switch technology to detect miRNA in the CFS and to incorporate the exponential amplification reaction (EXPAR) to bring the detection to clinically relevant levels. We assessed various EXPAR DNA templates under different temperatures and additives to enhance the accuracy of the sensing platform. Furthermore, different structures of toehold switches were tested with either high-concentration synthetic miRNA or EXPAR product to assess sensitivity. Herein, we elucidated the mechanisms of the toehold switch and EXPAR, presented the findings of these optimizations, and discussed the potential benefits and drawbacks of their combined use.

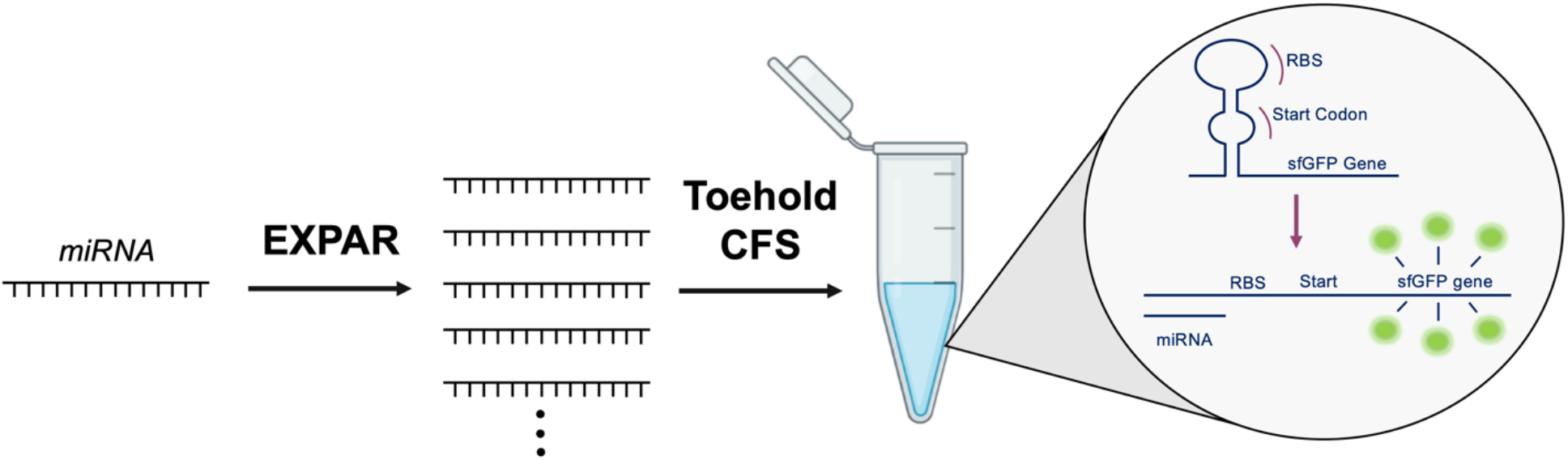

## 1. INTRODUCTION

MicroRNA (miRNA) has been highlighted as an important biomarker of various diseases. Recent studies have confirmed that the dysregulation of miRNA can be correlated with the development and specific stage of major diseases, multiple cancer types ^1-5^, viral infections ^6-9^, diabetes ^10-13^, and cardiovascular diseases ^14-16^. However, the current miRNA detection methods often require high-cost equipment and resources as well as suffer from low detection accuracy caused by miRNA vulnerability ^17-19^. Hence, there is a growing demand for new technologies for on-demand, low-cost miRNA sensors because of their potential to detect and monitor various diseases’ diagnoses and prognoses in non-laboratory settings. Most of the non-conventional low-cost methods rely on miRNA hybridization to an RNA or DNA in both the amplification and detecting modules ^17, 20^. This nucleic acid hybridization technique is the core concept of the toehold switch riboregulator, which has shown great success in detecting viral RNAs integrated with a cell-free system (CFS) ^21^. Although CFS’s portability and low-cost nature as a biosensor and this combination have the potential to be a robust and accurate miRNA detection platform, toehold switches detecting miRNA in the CFS have not been investigated previously.

The toehold switch is a programmable RNA sensor that detects the sequence of interest on the sample’s RNA called the trigger RNA. The synthetic riboregulators use toehold-mediated linear-linear RNA-RNA interactions designed *in vitro* that initiate strand displacement interactions ^22^. Once the trigger RNA binds to the target strand, the switch releases a sequestered ribosome binding site and a start codon to initiate the translation of reporter protein to verify if the RNA sequence of interest is present (Figure 1A). Nucleic acid sequence-based amplification (NASBA) is a pre-step isothermal amplification reaction ^23^ technique that is known to be fast and sensitive and has been used for diagnostics in field-based settings ^24^ as well as used with the toehold switch to bring femtomolar RNA concentration amounts in samples up to detectable levels ^21^. However, since typical 18-24 nucleotide long miRNAs do not comply with the length of the primer standards for the amplification, NASBA remains an unfeasible approach to amplify the miRNA up to a detectable amount that can operate the toehold switch ^21, 25^.

**Figure 1.**
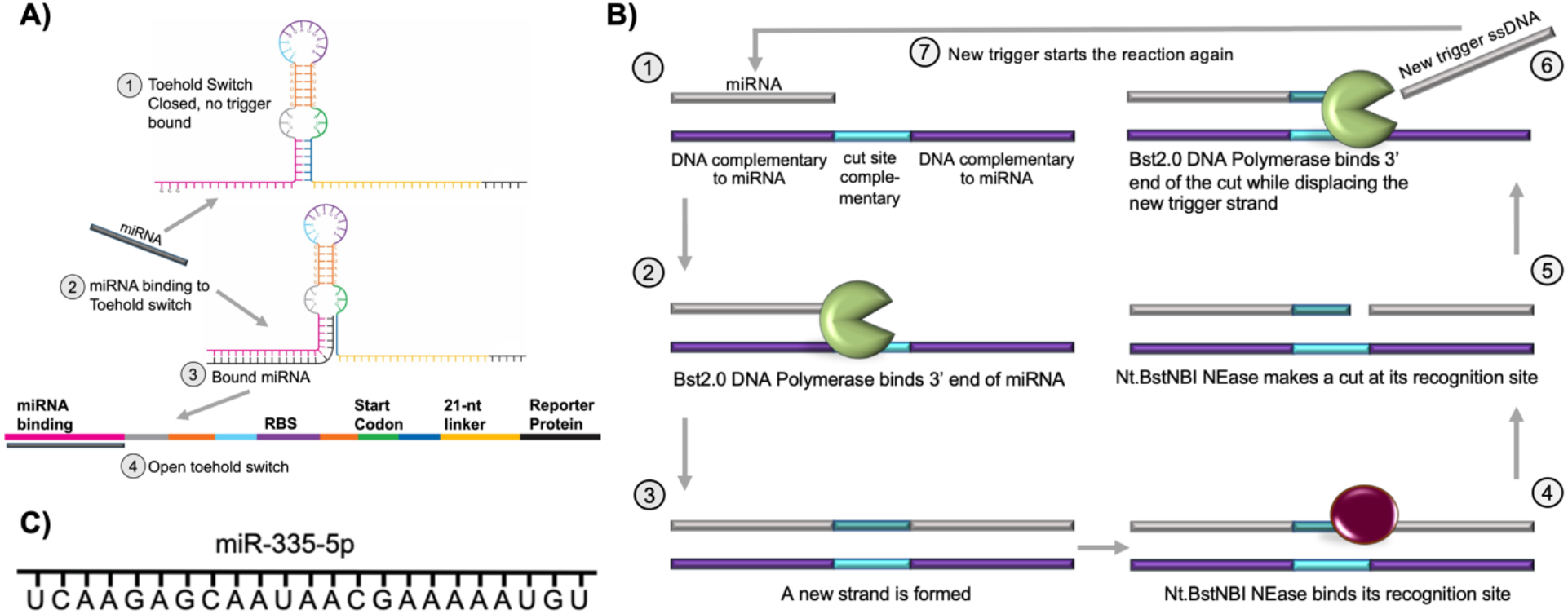
Combination of toehold switch and exponential amplification reaction for miRNA detection. **A)** Structure of toehold Switch. Once a complementary RNA molecule binds to the unsequestured region (step 3), it opens the switch by continued base pairing within the toehold (step 4). The opened toehold switch releases the start codon and ribosome binding site (RBS) to initiate reporter protein translation. **B)** Exponential Amplification Reaction (EXPAR) process. The trigger X (miRNA) anneals to the X’ region on the 3’ end of the template allowing for the 3’ end of the trigger to be exposed (step 1). The trigger itself acts as a primer, and a DNA polymerase binds to the 3’ end of the trigger extending the strand to the end of the other X’ region (steps 2-3). NEase creates a nick at the cut site (steps 4-5), causing DNA polymerase to come to the newly made 3’ end and displace the identical trigger strands (step 6). The displaced strand becomes a new trigger which can then start another reaction, and the initial trigger can start the reaction again on the same template strand (steps 7-1). **C)** Sequence of synthetic miRNA-355-5p.

On the other hand, exponential amplification reaction (EXPAR) facilitates isothermal miRNA amplification using short DNA oligonucleotides with half of the oligo binding to the target miRNA and the other half serving as a template to create multiple copies once triggered. This method can reach high amplification of >10^6^-fold in a short period of time (<30 minutes) ^26^. The reaction relies on a linear template containing two repeat regions (X’-X’) that are complementary to the trigger (target) sequence (X). These regions are separated by a short nicking endonuclease (NEase) recognition region. First, the trigger X anneals to the X’ region on the 3’ end of the template allowing for the 3’ end of the trigger to be exposed. The trigger itself acts as a primer, and a DNA polymerase binds to the 3’ end of the trigger extending the strand to the end of the other X’ region. Then, the NEase creates a nick at the cleavage site, causing DNA polymerase to start displacing identical trigger strands on the newly made 3’ end (Figure 1B) ^26, 27^. Despite the advantage of amplification of trigger strands, EXPAR suffers from the background amplification ^28^. Niemz *et al*. showed that the EXPAR template itself was one of the main generators of the background noise, and when the template sequence was changed, the background changed simultaneously ^28, 29^. Two main contributors of background amplification were the extension of the 3’ end due to intramolecular or intermolecular template-template non-specific binding and non-specific binding in general ^26^.

Here, we combined the EXPAR and the toehold switch in the CFS to develop the miRNA detection biosensor. We studied methods of reducing background amplification from EXPAR by comparing five different modified DNA templates, reaction temperature, and crowding agents. After optimization of amplification, the reaction showed the largest reduction of the background was chosen to be introduced to the toehold switch CFS compared to a high concentration of synthetic miRNA (miR-335-5p) related to tumor suppression as a model biomarker (Figure 1C). Two versions of toehold switches were tested with varying sequences to optimize the ratio of leak to ease of triggering. With these combinations, a robust, low-cost, and accurate miRNA detection system is expected to be created. Throughout the studies, we observed the limitation of toehold switch operation by miRNA trigger. We also became aware of how the miRNA interacts with synthetic sensing systems and the difficulties in detecting miRNA utilizing a toehold switch. However, the discoveries on the abilities of the EXPAR amplification system and the toehold switch sensing platform in the CFS were found and important to share for future research on these topics.

## 1. RESULTS AND DISCUSSION

### 2.1. Template Type and Temperature Study for Reducing Background

Biotinylation and phosphorothioate modifications of the EXPAR DNA template are the widely accepted methods to reduce background amplification. Biotinylated EXPAR templates reduce the melting temperature (*T*_*m*_) of the primer/template duplex ^30-32^. The 5’ end modification allows the primer to bind to the correct side of the EXPAR template (the 3’ side), and double modifications to both 5’ and 3’ ends help avoid non-specific binding. In addition, streptavidin assists decreasing the *Tm* of the duplex down 10 °C ^31^. It is believed that this decrease in Tm is due to steric repulsion between the protein and the binding site of the major groove of the DNA helix ^31^. Protection of DNA template degradation has also been found with the addition of biotin and streptavidin ^32^. Phosphorothioate modifications to the 5’ end of the template have shown promising results in reducing background amplification, mainly by protecting the DNA template from degradation ^33-38^. A recent study has discovered that bacterial post-transcriptional phosphorothioate modification positioned at a designated cleavage site or next to one can prevent digestion by the endonuclease ^39^. Likewise, the synthetic phosphorothioate internucleotide linkages enhance nuclease tolerance by replacing non-bridging oxygen in the DNA sugar-phosphate backbone with a sulfur atom ^40^.

Five different non-modified and modified (biotinylation and phosphorothioate) EXPAR templates were tested to find out if the template modification had a significant impact on reducing background amplification. We also investigated the reaction temperatures in streptavidin-supplemented amplification condition with the biotinylated templates (T4 and T5), as well as the range of reaction temperature to find the optimal condition of the lowest background, earliest detection, greatest sensitivity, and highest amplification (Figure 2). The templates include a standard EXPAR template (T1), three consecutive phosphorothioate bonds on the 3’ end only (T2), three consecutive phosphorothioate bonds on the 3’ end and three on the 5’ end (T3), a biotinylated tag on the 5’ end only (T4), and biotinylated tags on the 3’ and 5’ end (T5) (Figure 2A). T2 was the only template that had a significant time delay between the true signal output and background amplification, with a 15.2 and 12.3 minutes delay between 1 nM and 0.1 nM of synthetic miRNA, respectively (Figure 2B). However, reactions with all templates, including T2, showed high variance in amplification time of the true signal output and the background signal, making it difficult to determine when to stop the amplification reaction to avoid the potential background signal generation. For example, in the T2 reactions, even though the background is on average 12-15 minutes away from the true signal, the time between a delayed true signal and an early false signal is 5.6 and 2.1 minutes for 1 nM and 0.1 nM of synthetic miRNA, respectively. When streptavidin was added to bind to the biotinylated templates (T4 and T5) to reduce the melting temperature of the oligo regions, there still was not a noticeable time difference of amplification between the true sample (1 nM miRNA) and background (0 mM miRNA) (Figure 2C). Finally, we observed a large reduction of the background signal for templates T1, T2, and T5 at 53.9 °C, which was 1.1 °C lower than the standard reaction temperature (55 °C) (Figure 2D). This phenomenon could be due to the reduction of temperature, allowing easier complete binding of the synthetic miRNA.

**Figure 2.**
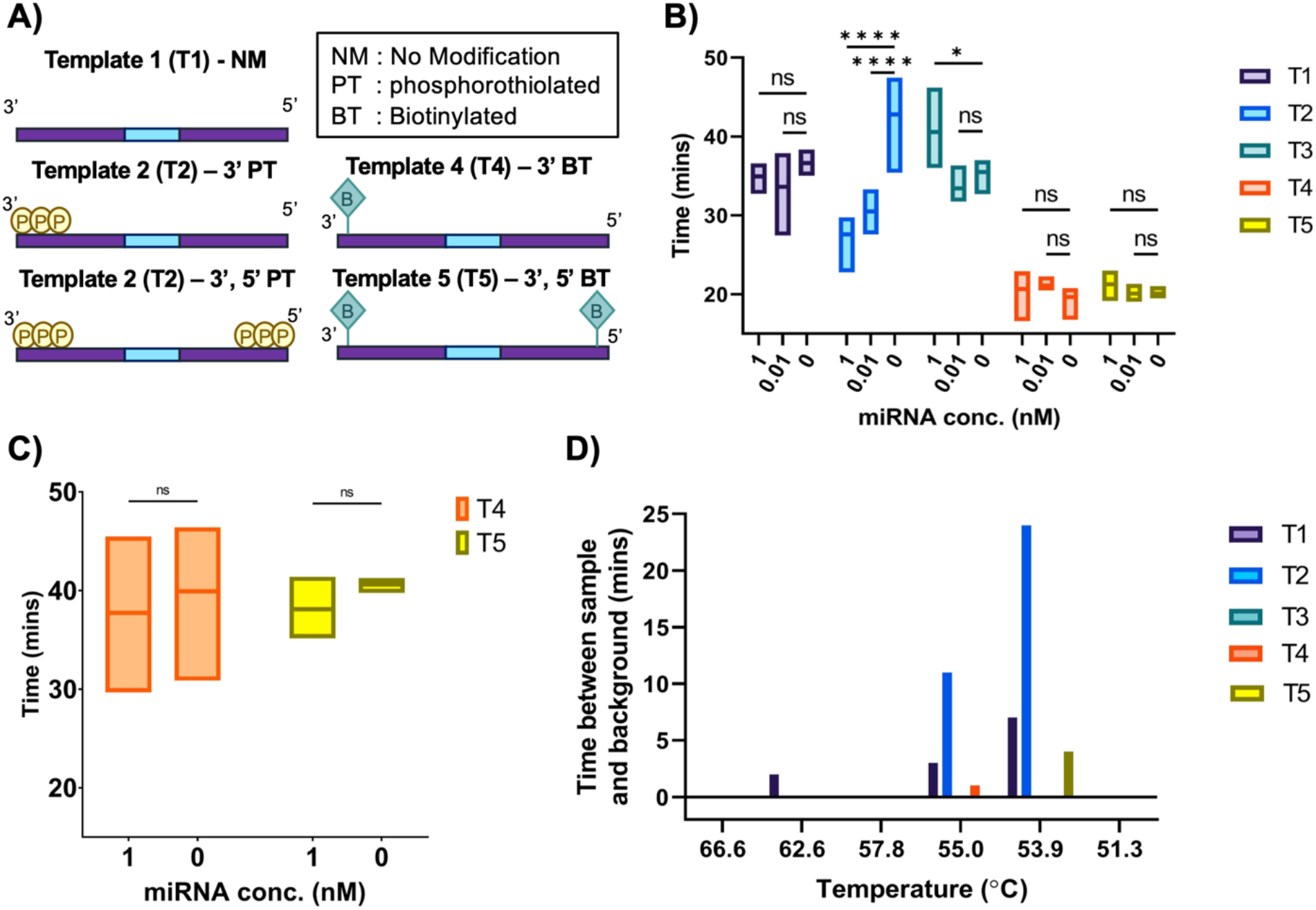
EXPAR template types and temperature comparison. **A)** Five EXPAR template modifications. **B)** Measuring background delay depending on miRNA concentration and template type. Values represented as mean ± SD, n=3, ****p<0.0001, *p<0.5. **C)** Addition of streptavidin to the biotinylated templates showed no significant delay of background amplification **D)** Effects of background delay depending on temperature and template showing more delay.

### 2.2. Molecular Crowding and Single-Strand Binding Protein

Macromolecules occupy 20-30% of the cells’ interiors and macromolecule crowding generates thermodynamic and kinetic effects on the cell’s functions ^41, 42^.When a spherical macromolecule occupies a space, the effective volume of occupancy is 8 times its intrinsic volume because it excludes other molecules from taking up that space and this increases thermodynamic activity. Brownian motion creates unsymmetrical forces from the contact of molecules on molecules ^43^. These forces increase the volume available to the macromolecules and satisfy the second law of thermodynamics by favoring maximum entropy. Molecular interactions under entropic forces could be more readily reversible than those caused by ionic forces (e.g., DNA condensed by polyethylene glycol (PEG) is more flexible and less compacted than when it is condensed by electrostatic interactions), thus encouraging the cell to correct mistakes ^43^. Single-strand binding protein (SSB) is a widely used protein that binds to single-stranded DNA (ssDNA). SSB can not only protect the unpaired ssDNA from degradation but inhibit secondary structure formation ^26^.

The crowding effect was tested by introducing PEG (2%) and SSB (2 μM) into the EXPAR reaction. The EXPAR fluorescence output was measured at minute intervals using a qPCR machine. First, we found PEG allowed for a slight reduction of background signal in the T5 template compared to 0% crowding, but not a significant difference compared to the present (1 nM) and absent (0 nM) of the synthetic miRNA (Figure 3A). In a separate experiment comparison, the amplification results of the combination of PEG and T5 were inconsistent, showing large variability across experiments with EXPAR in general that has been observed (Figure 3B). However, 2% of PEG induced a modest reduction of the background signal with the T2 template (Figure 3B). Unfortunately, we observed successive result inconsistencies when comparing the results of another EXPAR reaction (Figure 3C). This trend of inconsistency with the ability to differentiate low concentrations of miRNA from background amplification is an unattractive characteristic of EXPAR for miRNA sensing and amplification use. The substantial increase (x1000 times, 1 nM to 1 μM) of miRNA concentration input induced a noticeable signal output time difference with the T2 template (Figure 3C). However, this concentration (1 μM) is not clinically relevant. In addition, using this large concentration of miRNA did allow for a closer look at the crowding effect on the reaction, with SSB addition exhibiting a significant decrease in the background whereas PEG increasing background amplification (Figure 3C).

**Figure 3.**
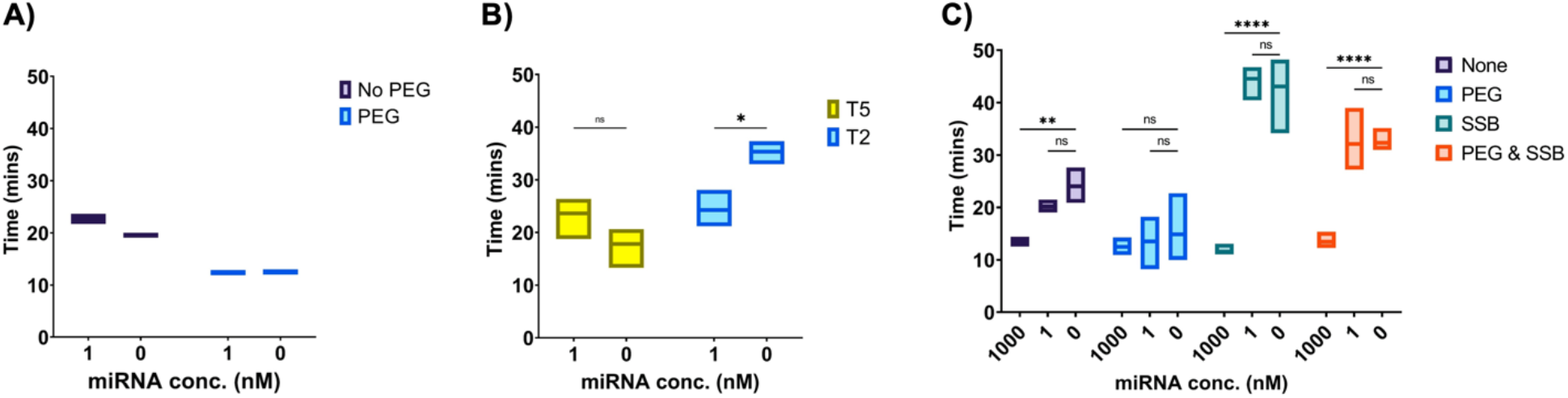
Effect of crowding agent and single-strand binding protein. **A)** T5 biotin template background reduction with and without crowding reagent (PEG). **B)** Reduced background under crowding conditions with biotinylated T2 and T5 templates. **C)** Combination of PEG and SSB on excessive miRNA concentration with T2 template. Values represented as mean ± SD, n=3,****p<0.0001, **p<0.01, *p<0.5.

### 2.3. Failed Toehold Switch Reactions and miRNA Hybridization Binding Energy

Once the miRNA is amplified from a low clinically relevant level, it should be able to be detected in the CFS miRNA sensing platform by triggering the toehold switch, giving a colorimetric output. Pardee et al. used thermodynamic considerations to optimize the toehold switch design by reducing the ribosome binding site contained loop sequence, further stabilizing the stem sequence of the sensor by adding extra nucleotides and removing the downstream refolding domain. In contrast to 18-24 nt short miRNAs, target-specific toehold switch optimization is a rational process with relatively lengthier target viral RNA strands. These factors have a large impact on decreasing leakage of false ON states (false positive) while increasing the free energy required to unwind the sensor ^21^, making it more of a challenge to tailor the toehold switch for miRNA triggers. Overall, their optimal toehold switch design contains a leader sequence of three guanines to encourage efficient transcription by T7 RNA polymerase, the exposed complementary trigger section (around 12 nt), a sequestered 9 nt continuation of the complementary sequence to the trigger, a conserved sequence region which contains the sequestered start codon in a 3 nt bulge and the sequestered RBS in a 12 nt loop, an exposed 21 nt low-weight amino acid coding sequences, and the protein of interest sequence ^21, 22^.

Based on the optimal parameters of the toehold switch, two versions (V1 and V2) of the toehold switch were designed since short miRNAs are required additional sequence spacers at the trigger binding site. Toehold switch V2, in particular, the loop length was elongated from 11 nt to 24 nt, which was shown to have the most considerable effect on increasing signal while keeping false triggering remains low. CFS reactions were carried out using 100 μM of synthetic miRNA, EXPAR product, or EXPAR background amplification product. The true EXPAR product was taken from the previously optimized reaction: 1 μM of miRNA, three consecutive phosphorothioate modifications on the 3’ end only (T2), and SSB. The reaction was then stopped after the expected true amplification detection and before generating the background amplification. The EXPAR background amplification product was taken from the EXPAR reaction with the same characteristics minus the miRNA trigger. It was allowed to run for 60 minutes for background amplification to occur. CFS reaction controls included miRNA or EXPAR input without toehold switch DNA, thus no fluorescence from the CFS should occur. Unfortunately, neither the synthetic 100 μM miRNA trigger nor the EXPAR product was able to trigger either version of the toehold switch, as shown by the lack of fluorescent change compared to the controls (Figure 4).

**Figure 4.**
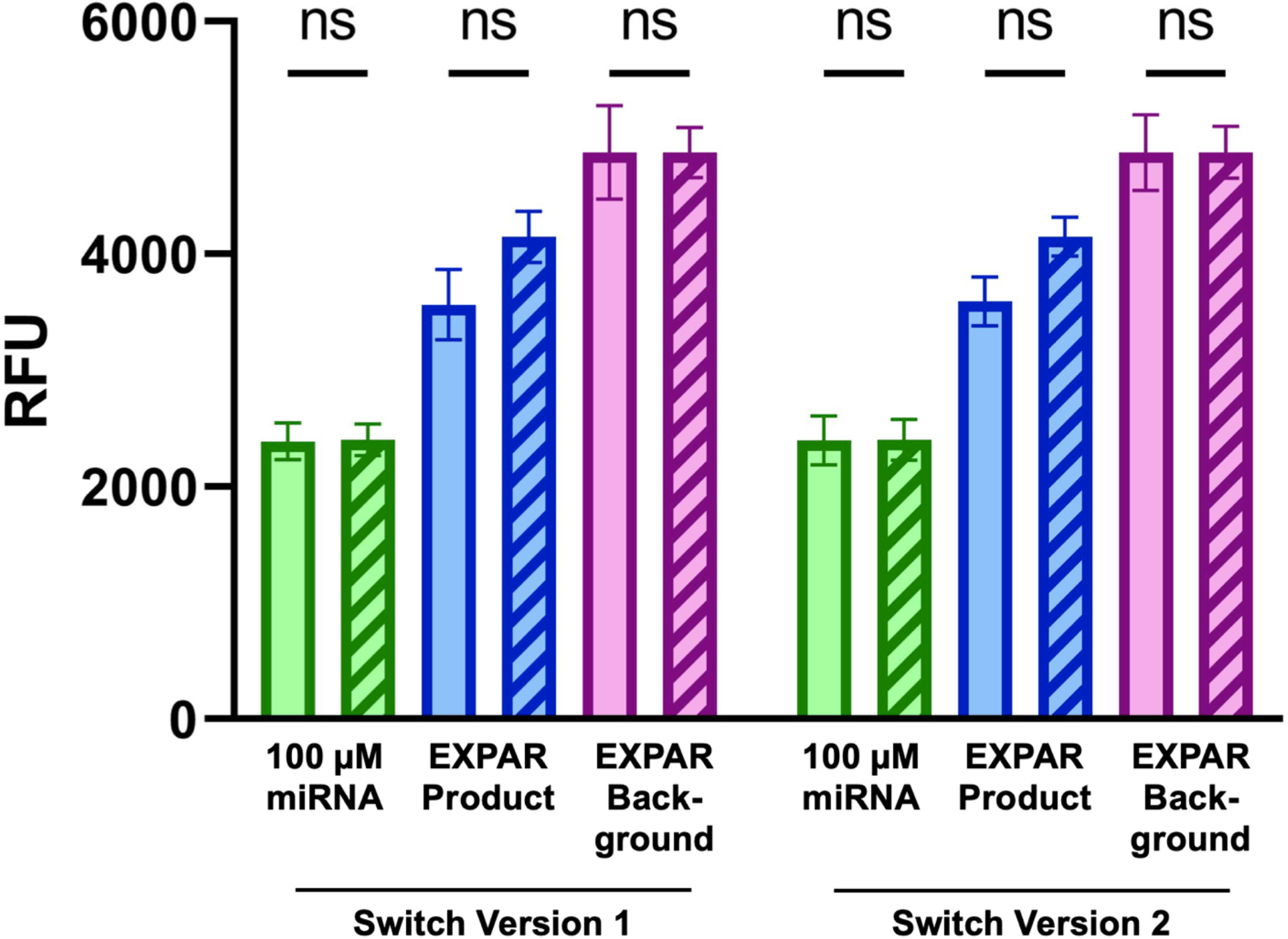
Toehold switch performance in the CFS. (Solid bars) Cell-free reaction with the synthetic miRNA at high concentration (100 μM), EXPAR product, or EXPAR background amplification product. (Pattern bars) Control reactions: Cell-free reaction containing no toehold DNA with no reporter fluorescent protein gene. Values are represented as mean ± SD.

## 3. CONCLUSION

The intrinsic open nature of the CFS allows for transforming the system into a versatile in vitro biomarker-sensing platform. In combination with the EXPAR of trigger miRNA sequence and programmable toehold switch, the CFS potentially can accomplish rapid, low-cost, accessible, and accurate miRNA detection system. However, due to the short length and low GC contents of matured miRNA, once bound to the toehold switch trigger, there are not enough nucleotides in the unbound state to attribute to base-pair stacking and other thermodynamics of the nucleic acid strand to counteract the repulsion caused by the negatively charged phosphate backbone ^44-46^. This results in weak hybridization to the binding region of the toehold switch and a lack of energy to operate the toehold switch.

The same principles can be used to account for EXPAR’s pitfalls, except for weak or incomplete binding causes false activation. miRNA, EXPAR products, or other free-floating EXPAR templates can bind incompletely to the binding region of the EXPAR template, causing an activation of the DNA polymerase even if the binding event is short-lived. Once the DNA polymerase is activated and displaces the complementary sequence to the binding region, the EXPAR product will be increased by 2^n^. Since this inaccurate EXPAR triggering is a random event, the reaction termination before the background signal generation cannot be relied on because it can occur at different times for different individual runs. There is also no alternative method to differentiate the background signal from the true sample signal output by end-point quantification using a qPCR machine, since they both will occur until the reaction runs to completion once triggered. The reaction conditions can only be reliable if the background amplification is removed completely.

In order to accomplish miRNA sensing, various new ideas have been suggested in the past few years, including isothermal amplification, electrochemical methods, enzymatic and other biochemical reactions, and nanoparticles ^47-49^. Unfortunately, the aforementioned approaches also resulted in relatively higher false positive rates ^19^, and the problem of inadequate specificity still exists. Specificity is the key to making an accurate diagnosis or prognosis of various diseases by miRNA detection due to high homology across miRNA sequences ^47^. However, many of the existing miRNA detection methods heavily rely on base-pairing of the short, single-stranded, mature miRNAs. Such hybridization methods lead to problems where a probe only partly binds to a miRNA, making it difficult to determine the accuracy of a putative signal ^47^.

In conclusion, utilizing EXPAR and the toehold switch in the CFS for miRNA sensing was not considered an optimal combination for the development of an accurate miRNA detection platform. Nonetheless, this study proposed a preliminary idea and framework to navigate the research guideline for designing miRNA detection platforms.

## 4. MATERIALS AND METHODS

### 4.1. In silico design

The toehold switches were designed *in silico* to detect miR-335-5p, with similar characteristics to the toehold switch series B design from Pardee *et al*. ^21^. Using NUPACK software, the optimal free energy and any conflicts of the secondary structure were found (Figure S2) ^50^. Sequences were constructed to be complementary to the miRNA target, form the toehold switch in the correct fashion desired, and ensure there were no stop codons or frameshifts after the start codon in the linker region. EXPAR templates were designed to include a complementary binding region, a nicking region, and another region for the polymerase to make identical copies of the miRNA in double-stranded DNA form. The genes were synthesized from Integrated DNA Technologies (IDT, Coralville, IA) with different template modifications (biotinylated and phosphorothioate additions) chosen in the settings of the IDT’s online builder. Synthetic single-stranded miRNA-335-5p was also synthesized from IDT.

### 4.2. Exponential Amplification Reaction (EXPAR)

EXPAR was conducted as specified by Zhang *et al*. ^51^ with slightly modified reaction components. The reaction was 20 μL in total, comprising of Mixture A and Mixture B. Mixture A contained trigger (synthetic single-stranded miRNA-335-5p), 0.2 μM amplification template, 1x NEBuffer 3.1, and 500 μM dNTPs. Mixure B contains 1x IsothermoPol buffer, 0.5 U/μL Nt.BstNBI nicking endonuclease, 0.5 U/μL Bst DNA polymerase, 2x EvaGreen^®^, and 2x ROX reference dye. Mixture A and Mixture B were prepared separately on ice, and then they were pre-heated separately to 55 °C before combining Mixture B into Mixture A. The fluorescence output was measured at a minute interval using the qPCR machine (Bio-Rad, Hercules, CA) immediately. The qPCR machine was set to run at 55 °C for 70 cycles. Streptavidin was added to Mixture A containing the biotinylated templates with a final concentration of 2 μM in Mixture A+B. Single-strand binding protein from Sigma-Aldrich (S3917) was added at the final concentration (in Mixture A+B reaction) of 2 μM to Mixture A as described ^26^. Polyethylene Glycol (PEG 8000, Thermo Fisher Scientific, Waltham, MA) was added to Mixture A at a final concentration of 2% in the Mix A+B reaction. All enzymes and ROX reference dye were purchased from New England Biolabs (Ipswich, MA).

### 4.3. Recombinant plasmid

The toehold switch was added to the pJL1-*sfGFP* vector following the standard Gibson Assembly method. The recombinant plasmid was then transformed into DH5alpha *Escherichia coli* competent cells via electroporation and selected on an LB agar plate containing Kanamycin (50 μg/mL). Colonies were screened by colony PCR, and subsequent Sanger sequencing confirmed the sequences. The recombinant plasmid was then purified at a high concentration (200 μg/mL) using the plasmid preparation kit (Qiagen, Germantown, MD).

### 4.4. Cell-Extract Preparation

The *E. coli* strain BL21 Star™ (DE3) (genotype F-*ompT hsd*S_B_ (r_B_-, m_B_-) *gal dcm rne131* (DE3)) (Invitrogen, Waltham, MA) was used for the cell-extract for the cell-free protein synthesis reaction. The cell-extract was prepared as described previously ^52^. In brief, the cells were grown in 2xYTPG media in shake flasks at 37 °C with vigorous shaking (250 rpm). The cells were harvested OD_600_ at 3.0 and washed three times with the chilled Buffer A (10 mM Tris-acetate (Ph 8.2), 14 mM magnesium acetate, 60 mM potassium acetate, and 2 mM DTT). The cells were then resuspended in Buffer A (without DTT) and lysed by sonication. The cell-extract was aliquoted out, flesh frozen in liquid nitrogen, and stored in -80 °C freezer until use.

### 4.5. Cell-Free Reactions with Toehold Switch

Cell-free reaction mixtures were prepared as described previously ^53^. The standard 15 μL cell-free reaction mixture in a 1.7 mL microtube includes BL21 Star™ (DE3) cell-extract (4 μL), 1 μL salt solution (12 mM magnesium glutamate, 10 mM ammonium glutamate, and 130 mM potassium glutamate), 1.2 mM ATP, 0.85 mM each of GTP, UTP, and CTP, 34 μg/mL L-5-formyl-5,6,7,8-tetrahydrofolic acid (folinic acid), 170 μg/mL *E. coli* total tRNA, 57 mM HEPES buffer (pH 7.2), 0.4 mM nicotinamide adenine dinucleotide (NAD), 0.27 mM coenzyme A, 4 mM sodium oxalate, 1 mM putrescine, 1.5 mM spermidine, 2 mM each of 20 canonical amino acids, 33 mM phosphoenolpyruvate (PEP), and 13.3 μg/mL plasmid. The cell-free reaction microtubes were then incubated at 37 °C for 20 hours. Synthetic single-stranded miRNA-335-5p, EXPAR product from miRNA, or EXPAR background product from no miRNA present were inserted into the cell-free reaction with the toehold switch plasmid. Negative control reactions included the cell-free reaction mixture with either synthetic single-stranded miRNA, EXPAR product, or EXPAR background product with no plasmid DNA containing the toehold switch and sfGFP gene.

### 4.6. Statistical analysis

Statistical analyses were conducted using GraphPad Prism 8.4.3 (GraphPad Software) with a 5% significance level. For the parametric analysis of data from the fluorescent measurements from synthesized sfGFP, two-way ANOVA was used.

## NOTES

The authors declare no competing financial interest.

## ACKNOWLEDGEMENTS

This work was supported by the LSU LIFT^2^ Fund (Grant No. AG-2022-LIFT-002) and the USDA National Institute of Food and Agriculture HATCH fund (Accession No. 1021535, Project No. LAB94414).

