## Supporting Information for "Suitability Evaluation of Toehold Switch and EXPAR for Cell-Free MicroRNA Biosensor Development"

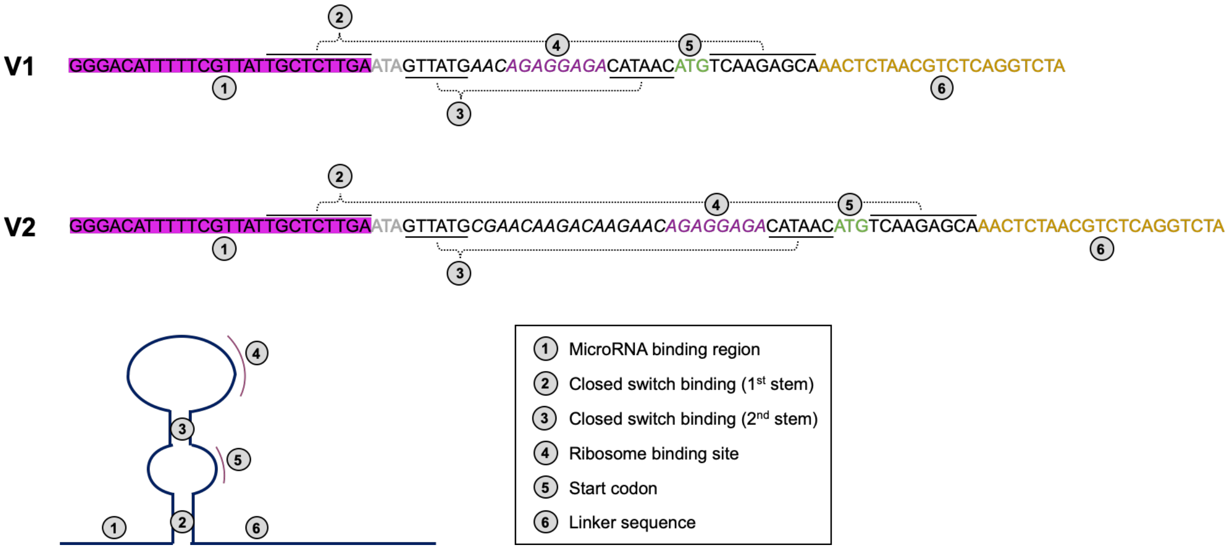


**Figure S1. Toehold switch designs (5' to 3') used in this study.**

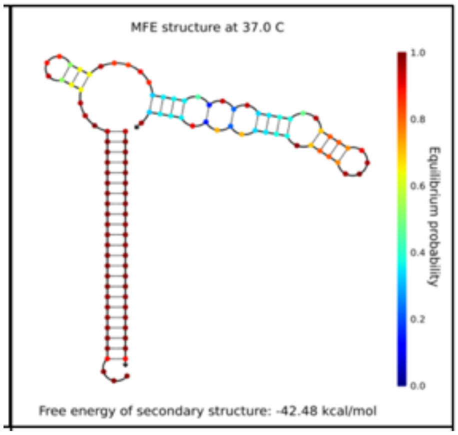


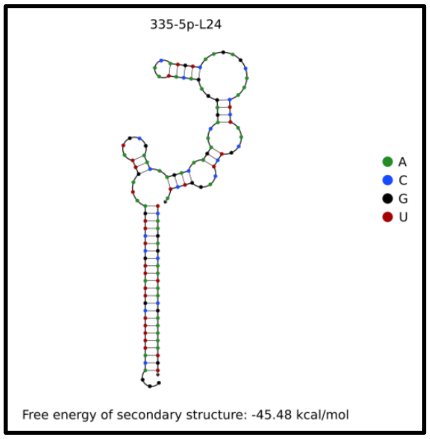


**Figure S2. NUPACK free energy analysis of the toehold switch V1 (left) and V2 (right) in complex with miRNA-335-5p**

**Table S1. EXPAR template sequences (5' to 3') used in this study.**

| Template | Sequence (5' - 3') |
|---|---|
| T1 | ACATTTTTCGTTATTGCTCTTGATTCAGACTCAAACATTTTTCGTTATTGCTCTTGA |
| T2 | ACATTTTTCGTTATTGCTCTTGATTCAGACTCAAACATTTTTCGTTATTGCTCT*T*G*A |
| T3 | A*C*A*TTTTTCGTTATTGCTCTTGATTCAGACTCAAACATTTTTCGTTATTGCTCT*T*G*A |
| T4 | A/iBiodT/CATTTTTCGTTATTGCTCTTGATTCAGACTCAAACATTTTTCGTTATTGCTCTTGA |
| T5 | A/iBiodT/CATTTTTCGTTATTGCTCTTGATTCAGACTCAAACATTTTTCGTTATTGCTCTTG/iBiodT/A |

Underlined: Nicking endonuclease recognition sequence

*: Phosphorothioate modification

/iBiodT/: Internal biotin tag
